# Simulating the Idling Behavior of Escherichia Coli

**DOI:** 10.1101/189878

**Authors:** Herbert Weidner

**Author notes:** 16. September 2017.

## Abstract

Until now, it is not known how the motors of E. coli reverse the direction of rotation, mostly simultaneously. If the C-ring is bistable and if the number of bound CheY-P depends on the direction of rotation, there is a strong variation in the concentration of CheY-P in the cytoplasm. Then, the synchronous changeover can be explained without further additional assumptions.

## Introduction

E. coli bacteria react to external stimuli, more specifically, to concentration gradients of chemical attractants or repellents. In order to find more suitable locations, the rotation direction of the drive motors is reversed at irregular intervals, which is why each bacterium alternately either tumbles (as long as the flagella rotates clockwise) or swims forward (the flagella rotates CCW). Runs are the default behavior, with intermittent tumbles. Since this genetically fixed behavior does not stop in a chemically constant environment, we call it *idling behavior*. As the concentration of food or repellents changes, the duration of runs and tumbles changes slightly, and the bacterium moves slowly on a zigzag course into a region with better survival conditions. Obviously, the tiny bacterium is able to produce a proper reaction by comparing the current sensory data with historical data.

One could learn a lot about the function of the bio-computer in E. coli if it were possible to reproduce the behavior on an electronic computer. But clocks, memory and comparator circuits have not been found in any bacterium. Bacteria perform the task with the help of several thousand signal proteins such as CheY. Every molecule can change its location and its neighbors, some molecules can change their shape or connect with others during a short period. It would be quite interesting to understand how the chaotic "dance of molecules" can produce meaningful behavior.

Perhaps, *reverse engineering* can help to understand how the bio-computer supports a bacterium in the search for food. We can observe how these bacteria behave. What happens internally to create their behavior? We know that the software of E. coli deviates strongly from computer software. The genes describe only the structure of the proteins but do not contain any rules of behavior. The structure allows or prohibits which molecules can connect as soon as they touch.

We can read the genetic code, but do not understand it. As with an unknown language, we must first learn some basic facts. It is perhaps a good idea to start with a bottom-up approach and try to simulate the only observable behavior of the bacteria, the incessant and irregular reversals of the rotation direction of the motors. In order to investigate this “idling behavior”, we ignore all variable external influences and restrict to the known properties of the biochemical building blocks in the cytoplasm and the known methods of technical control technology.

The successful simulation of the idle activity is a prerequisite for a promising study of how external signals influence the behavior of the bacteria.

> You don’t understand it until you can simulate it.

## Deciphering the idle mode of the engines

Living E. coli bacteria are never quiet. This means a waste of energy in the drive motors, because when all sensors of a bacterium report constant signal strength, there is no need to leave the current location. Nevertheless, no bacterium ceases to move on a zigzag course and remains within a narrow range around its current position. Why? Little helpful seems a reasoning like: ”¨it is desirable for the cell to resume normal behavior continuing the search for better conditions (e.g. nutrients)." Can bacteria actually decide between different behaviors?

We do not believe that E. coli consciously optimizes its behavior, we suspect a rather automatic process, a technically comprehensible solution, which should be explained on a molecular basis. Our goal is to model the idling behavior of E. coli while omitting variable external signals. Therefore, the simulation requires no modeling of the sensor complex.

As long as the attractant and repellent concentrations in the extracellular space is constant, the sensor complex at one end of the elongated cell is believed to produce a constant concentration of CheY-P in the cytoplasm. The signal molecules slowly diffuse to the motors and generate their idle behavior. In the case of technical signal transmission, the CheY-P would be dephosphorylated near the motors because the signal generated by the sensor complex has "arrived" at the target location and can be erased. This principle is presumably followed in B. subtilis, but not in E. coli. Here, the CheY-P either spontaneously decay after a several seconds or they diffuse back to the sensor complex, where they are destroyed by CheZ. (In the course of the simulation, we will see that this "feedback channel" could serve to report the motor activity to the sensor complex.)

From the point of view of information technology, it is puzzling how the motors can change their direction of rotation abruptly: The sensor complex is far away from the drive units and all information is transmitted by the slow diffusion rate of the signal molecules CheY-P. Since the bandwidth (the amount of information per time unit) is very low, the central information processing near the signal receptors can not suddenly reverse the rotation direction of the far-away motors. But this is exactly what is observed.

Hitherto it has been regularly assumed that the sudden reversal of the direction of rotation of the motors is somehow caused by molecular noise. This can not be true, because then both rotation directions should occur equally often and with similar mean duration. If noise were the cause, the synchronous switching of multiple motors would rarely occur - quite in contrast to the observation.

A rapid reversal of the rotation direction can only be effected by the motors themselves (or something in the immediate vicinity). The simplest explanation: Each flagellar motor is bistable and can change its structure very quickly thanks to a positive feedback like a Schmitt trigger. This known electronic circuit converts an analog input signal to a digital output signal. Even if the input changes very slowly, the changeover takes place very quickly after a certain threshold value is reached. A small hysteresis prevents unwanted frequent switching[^1^].

The flip-flop effect of the flagellar engine is the result of allosteric mechanisms of certain assemblies in collaboration with CheY-P molecules[^2^]. Using a single postulate, we explain - on a molecular level - how the CW-CCW switch of a flagellar engine may work:

At least one type of protein (FliM?) must recognize the direction in which a motor is rotating. This information must affect the number of CheY-Ps that the C-ring of the engine can bind.

Assumptions: Each flagellar motor in E. coli can switch between two different configurations A and B[^3^]. The flagellar switch protein FliM can connect to CheY-P, but not to CheY.

State A: The flagellar motor rotates counterclockwise (CCW) and E. coli runs. The docking site for the signaling protein CheY-P is easily accessible from the cytoplasm. As the number of CheY-P in the cytoplasm rises and / or the time progresses, more and more CheY-Ps are bound to FliM. As soon as a sufficient number of CheY-Ps are mounted, a certain upper threshold A (say 28) is exceeded and the configuration of the motor changes abruptly (allosteric switch)[^4^][^5^].
State B: Now, the motor turns clockwise and E. coli tumbles. In this state of the engine, the signal molecules CheY-P can be poorly bound and dissolve rapidly from the C-ring. That continues until too few CheY-Ps are coupled to the motor. As soon as the number of bound CheY-P decreases below the threshold B (say 4), the motor suddenly switches back to state A.

There are experimental indications that the FliM molecules can bind a different number of CheY-Ps depending on the direction of rotation of the motor[^6^]. Possible causes are steric hindrance due to displaced neighbors or allosteric deformation of FliM. This simulation does not deal with these molecular details.

The switch complex at the bottom of a flagellar motor is constructed from about 34 FliM proteins [^7^]. The successive addition of CheY-P deforms the C-ring elastically, it stores more and more energy. After a certain limit has been exceeded, the motor changes its shape like a snap-switch mechanism. It has to be clarified how strong this event reduces the ability of the C-ring to keep hold of CheY-P molecules. It is also unclear how fast the already bound CheY-P escape to the cytoplasm. Finally, the minimum number of bound CheY-P would be interesting, below which the basal body switches back into its original form (lower threshold).

The difference between the two thresholds (hysteresis) and the concentration of CheY-P in the environment determines the switching frequency of the rotational direction. While it is quite clear how external signals influence the concentration[^8^], it is still unknown whether and how the two switching thresholds can be changed. One fact is certain: Since only few CheY-P molecules are involved, strong statistical fluctuations must be expected. This is precisely what is observed - the direction of rotation changes very irregularly.

The process described above is robust to changes in biochemical parameters, such as protein concentration and temperature. It is particularly important that it also works with *constant* and almost arbitrarily high CheY-P production rate in the sensor complex. The concentration in the surroundings of the motors fluctuates strongly - the consequences are discussed in more detail below. These are the minimum requirements for a practicable idle program. The sensor complex and the drive units do not have to work precisely and coordinated, and there are no strictly tolerated numerical specifications.

The process is intrinsically safe: Even if the production rate of CheY-P (in the far-away sensor complex) deviates significantly from the usual value, the motor will not continuously rotate CW or CCW. In a hypothesis with very high hill coefficients, such a deviation is catastrophic[^9^][^10^].

There are several possibilities to change the duration of states A and B in order to adapt to changing environmental conditions, a basic requirement of life[^11^]. This expansion of the idling behavior is not yet discussed here. As with a diesel engine: The analysis of the load behavior must wait for the idle to function properly.

## Synchronization of motors

When all motors turn counterclockwise (CCW), the filaments work together in a bundle that drives the cell steadily forward - the cell runs for about one second; when *at least one* motor turns clockwise (CW), the bundle flies apart, and the motion is highly erratic - the cell tumbles.

If the flagellar motors are autonomous, how are their filaments able to work synchronously in a bundle? As the bacteria often swim longer distances, there must be an easily understandable, internal mechanism, which ensures that all motors of the same cell reverse synchronously[^12^]. Measurements clearly show that the directional switching between two motors is highly coordinated[^13^][^14^].

Statistical fluctuations in conjunction with a very large hill coefficient can probably not produce this simultaneity. It is questionable how realistic the veto model[^15^] is because it does not provide any explanation on molecular level. In contrast, the above-described process provides a very simple and *verifiable* explanation: As long as a motor rotates in CW, almost all bound CheY-Ps dissolve rapidly from the C-ring and increase the concentration of the CheY-P in the surrounding cytoplasm. This change in concentration spreads out like a damped wave in the cytoplasm and affects neighboring motors, probably even the sensors. If the distance between the motors is significantly lower than the wavelength, the motors will synchronize. This is a well-known physical phenomenon[^16^].

We have a self-organizing and self-reinforcing process: The more motors participate, the greater gets the amplitude of the perturbation and the stronger the remaining motors are influenced. Due to the limited number of CheY-P in the cytoplasm, no instability can occur.

This statement can be quantified using the numerical values from the "digital model" described below. The E. coli cell is about 3 μm long and we assume a distance of 0.3 μm between two motors. In our model, we choose the box length 3.8 nm. So the wave must travel 80 box lengths, which requires approximately 1000 time steps (0.86 μs each, see below). At a phase speed of 350 μm/s, the wave reaches the adjacent engines within a few milliseconds. Since the observed operating periods of CW or CCW rotations last substantially longer, the described synchronization is easily possible. The simulation below shows that no additional assumptions are necessary. Synchronization of the motors is an emergent feature of the bistable process described above.

All that is missing is the experimental evidence that the CheY-P concentration in the cytoplasm and the reversal of the rotation direction fluctuate synchronously.[^17^].

## The modeling of the cell interior

E. coli is a 3 μm long <u>**cylinder**</u> with a diameter of 0.8 μm. We model it by a cuboid, consisting of 790*210*210 individual cubes (boxes) of edge length 3.8 nm, leading to a time step Δt = 0.86 μs (see below). It is not claimed that this decomposition is particularly close to reality, but right-angled volumes are computationally much faster and easier to handle than other shapes like cylinders.

In this model, we focus on the simulation of the idling behavior of the motors. We do not need any information about details of the sensor complex in E. coli, we need a single type of signaling molecule, CheY-P. Since no external signals are allowed, the sensor complex - far from the motors - ensures a constant number of these molecules in the cytoplasm. Other signal proteins such as CheZ are not required and are therefore ignored.

Inside an E. coli bacterium, 7100 CheY proteins[^18^] are distributed over a total of 4.5 million boxes. An unbound signal protein is never at rest; after each time step, it jumps into one of 26 surrounding boxes. Estimated 2% CheY are phosphorylated and diffuse through the cytoplasm, very few come close to a motor. Most boxes contain only other particles like H2O or ATP and are therefore ignored. In addition, five complex-built motors are available at freely selectable but fixed locations. We install the motors spaced about 0.3 μm apart. The simulation shows that the exact position of the motors hardly influences the results.

The engines (45 nm in diameter) are not point-like, they are represented by an immovable octagon of 32 boxes at the edge of the cytoplasm. Each box of this C-ring contains a single FliM mole-cule[^19^] and may bind one CheY-P. It is still unknown whether the signal molecules dock onto the inner or outer side of the C-ring. Due to lack of accurate data, a box size of 3.8 nm is appropriate.

During the simulation, it is counted how many CheY-Ps are bound to the FliMs of each engine. In the absence of more accurate data, we start with the upper threshold A = 28 and the lower threshold B = 4. The switching thresholds may possibly be fine-tuned by signal molecules other than CheY-P. During the simulation, these assignments may be adjusted in order to reproduce the observable idle behavior of the motors. Since the exact reason for switching the direction of rotation is still unknown, it is also unclear whether a special arrangement of the CheY-Ps on the C-ring is necessary.

Before starting the simulation, another problem must be solved at a molecular basis: according to the central assumption of this simulation, in state A, every FliM molecule can bind one CheY-P. As soon as the sum exceeds a certain threshold value, the C-ring switches to state B and most of the CheY-P disconnect within a short time. Do they stay in the immediate vicinity?

Probably not, because the motor is driven by a current of H^+^ ions (about 300,000 ions/s), entering the cell through the proton channels. This constant flux of the numerous charged particles produces a <u>**vortex ring**</u> in close proximity to the C-ring that drags the released CheY-P and disperses them quickly in the cytoplasm. Simplifying, we assume that this distribution starts at some distance from the engine in extension of the rotational axis in the so-called starting area. This mechanism prevents the released CheY-Ps from immediately re-attaching to the C-ring. The distance of the starting area from the engine influences the simulation result and must be examined more closely as soon as more engine details are known.

We do not discuss how the many protons are exported after they have fulfilled their task as "fuel" of the engines.

The most frequent and time-consuming task of the simulation is to check whether two signal proteins get in contact. In order to determine whether two of them get close enough to react with one another (= to transfer a signal), one does not have to know their exact locations. It is sufficient to find out whether they are in the same box. This speeds up the simulation significantly, because it is very easy and fast to check whether the coordinates of two particles match in a coarse grid. It would take much longer to calculate the true three-dimensional distance and to compare three floating numbers. Inside the cell, this knowledge is not very helpful, because each particle changes its place during a time step.

A simulation is supposed to imitate real behavior. Therefore, the program must be based on well-known natural laws and as little prejudice as possible should be installed. This may contradict the demand for simple programming and short computing times. It is hard to find a good compromise. In order to keep the calculations simple, we assume that the signal molecules move at constant speed and the *Brownian motion* of the neighboring molecules forces them to constantly change the direction. In this model, they jump into one of the 26 neighboring cells after each time step, but none may leave the bacterium. The direction is random because no signaling molecule can know that it must oscillate between the receptors and the propulsion motors to deliver messages. There is no physical reason for this assumption. Nevertheless, one finds exactly this unfounded assumption in previous simulations[^20^], probably to simplify the simulation and to shorten the computing time. Such arbitrary specifications render the results of the corresponding simulation worthless, because in this way, one can force any desired behavior.

## Connecting the digital model to the real world

There is a good reason to limit the number of allowed jump directions for molecules after each time step to only 26 neighbors: the permissible coordinates can be unambiguously identified by packed integer numbers, and in each computer, they can be processed much faster than floating-point numbers (see below: *Technical Details*). It must, however, be checked whether the diffusion of the molecules in the cytoplasm is correctly described despite this simplification.

It is mandatory to prove that the model can reproduce Fick's second law of diffusion without errors. This law helps to calculate how much time passes in the real world when the simulation performs one single step. The formula [^21^]

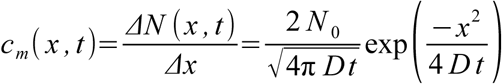

represents a relationship between the known value of the diffusion constant *D*[^22^] of the signal molecules in the cytoplasm of a cell[^23^], the time *t* and the spatial resolution *Δx* of the model. To check Fick's law, we regard the bacterium as a long tube and start with N_0_ = 7100 proteins near the left end. As the time progresses, they diffuse to the right but no protein must leave the cell. After 1000 steps of "Brownian motion", the fastest signal molecules have reached the box number 110. After 10,000 steps, the fastest proteins have reached the middle of the bacterium.

The picture shows the number of CheY proteins in thin slices (size of one slice = x/y/z = 1/210/210) transverse to the cylinder axis of the bacterium. The considerable statistical fluctuations of the local protein density (colored red and blue) are obvious. Since fractions of proteins do not fulfill their function, it is questionable whether smoothed, differentiable functions (black) permit a correct description of the reactions. Particularly in the vicinity of the motors, there are very few CheY-P - too few to calculate a serious statistic.

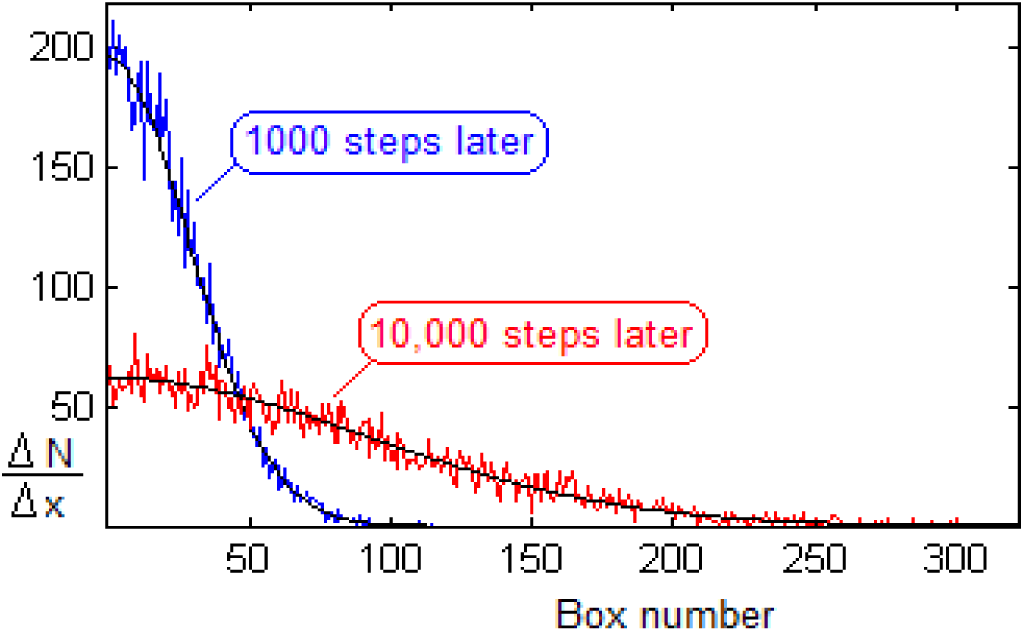

Inserting the model results for x = 0 into the formula above, we get:

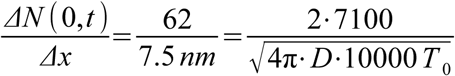

Assuming a diffusion constant *D* = 7 μm^2^/s[^24^][^25^] for the signal molecules CheY-P, the desired edge length of 3.8 nm for each box (see the discussion of the C-ring) forces the time step *T*_0_ = 0.86 μs.

The simulation of one minute real-time requires 70 million time steps. Depending on the quality of programming and the speed of the computer, the simulation may last longer.

## Choice of parameters

For the computer, nothing is self-evident. During the programming, a complete reaction must be prepared for every conceivable state, even if it seems very unlikely. For this reason, data must be given and decisions made which do not play any role in verbal discussions and are usually overlooked. A small selection:

- State A: Does the C-ring change its shape when more and more CheY-Ps are added? Does this affect the binding force between FliM and CheY-P? Could this cause some of the already attached CheY-Ps to be solved in order to symmetrize the C-ring?
- State B: How fast do the bound CheY-P separate from the FliM proteins and diffuse back into the cytoplasm? Is there any preferred arrangement or is it (in the extreme case) possible that one semicircle of the C-ring binds not a single CheY-P while the other is still full?
- When the C-ring changes its state (and the engine the direction of rotation), some CheY-P could be separated due to the "vibration". How big could the number be? The simulation shows that values between 0 and 5 do not produce any significant consequences.

1. Are the motors randomly arranged or is there a pattern? Are there minimum and maximum distances?

Chosen parameters: The C-ring consists of exactly 32 FliMs, threshold (A→B) = 28, threshold (B→A) = 4. In state A, the CheY-P adhere to the FliM of the the C-ring for an indefinite period of time and do not dissolve spontaneously. During state B, we choose a half-life T_½_ = 10 ms.

At the beginning of each simulation, the C-rings of all motors do not bind any CheY-P and the cytoplasm contains 156 CheY-P (= 2.2% of 7100 CheY). If the number is increased to 400 (6.5%), the duration of the longest run intervals drops below 100 ms and the bacterium tumbles mostly.

With even more CheY-P, E. coli tumbles almost continuously - quite contrary to all observations. The assumption[^26^] that 30% of all CheY are phosphorylated can not be simulated, because in the immediate vicinity of each motor, there are so many CheY-Ps that all 32 FliMs are occupied within a few milliseconds. Separated CheY-P are immediately replaced by others.

> You don’t understand it until you can simulate it.

In the model of size (790/210/210), the values (44/99/0), (133/209/124), (222/74/209), (311/97/209) and (400/209/139) were chosen as motor coordinates. The sensor complex would be near the other end (750 /xx /xx), it is not needed in this simulation.

The starting range is 114 nm (=30 boxes) apart from each motor. From this range, the CheY-Ps diffuse into the cytoplasm after they have separated from the C-ring after switching to state B.

## Simulation Results

The simulation of a single motor does not provide any great insights, the result corresponds to that of the known <u>**integrating ADC**</u> in electronics. The cytoplasm contains only 156 CheY-P, although each E. coli harbors approximately 7,000 CheY signaling molecules. This number was chosen to reach the experimentally observed time periods of "runs" and "tumbles". Since the C-ring can bind a considerable portion of this surprisingly small amount, strong, almost periodic changes in density occur in the cytoplasm. A decreasing amount of CheY-P reduces the frequency of directional switching because it takes longer to reload the C-ring. The frequency can also be changed by shifting the two thresholds or by changing the half-time in state B. Whether this fine tuning is the task of some puzzling signal proteins in E. coli has yet to be explored.

This simulation assumes that CheY-P is neither newly formed nor destroyed. Surely, the strong fluctuations of unbound CheY-P also spread to the signal complex, which will probably react. The nature and strength of this reaction is unclear and must be modeled separately - provided that the fluctuations in density can be confirmed experimentally.

In the next step, the simulation confirms what was theoretically foreseen: the motors influence each other because the cytoplasm contains only few signal molecules CheY-P. (This does not apply to the propellant protons, which are probably available in the extracellular space in an unlimited amount.)

In Fig. 4, an E. coli bacterium is covered with five motors distributed over the surface. In terms of programming, it has been ensured that there is *no* connection between the motors, except for a common supply of CheY-P.

**Fig. 1:**
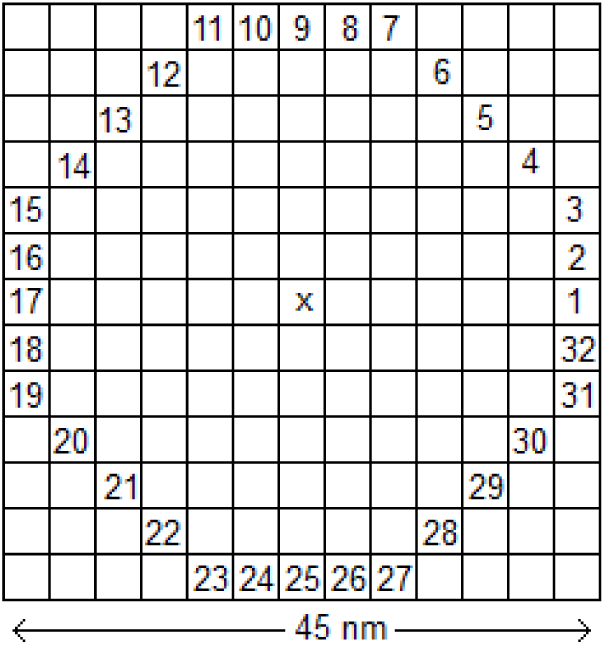
Top view of the C-ring of a motor in our model.

**Fig. 2:**
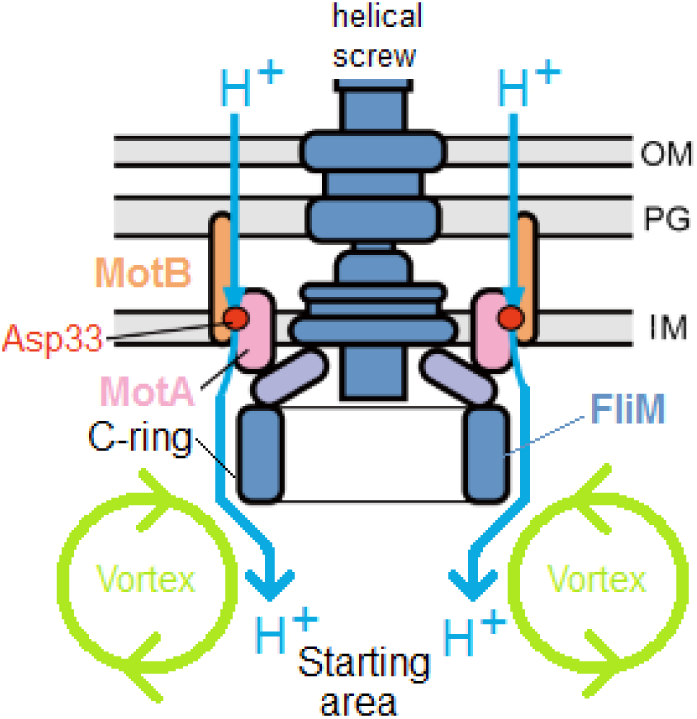
Sectional drawing of a drive motor with previously known assemblies (the C-ring has a diameter of 45 nm). The rotor rotates when large quantities of protons flow into the cell interior. If the direction of rotation is CW, the CheY-P are only weakly bound to the FliM. The proton current dissolves them and blows them into the cytoplasm. The resulting vortex causes the CheY-P to move away from the C-ring.

**Fig. 3:**
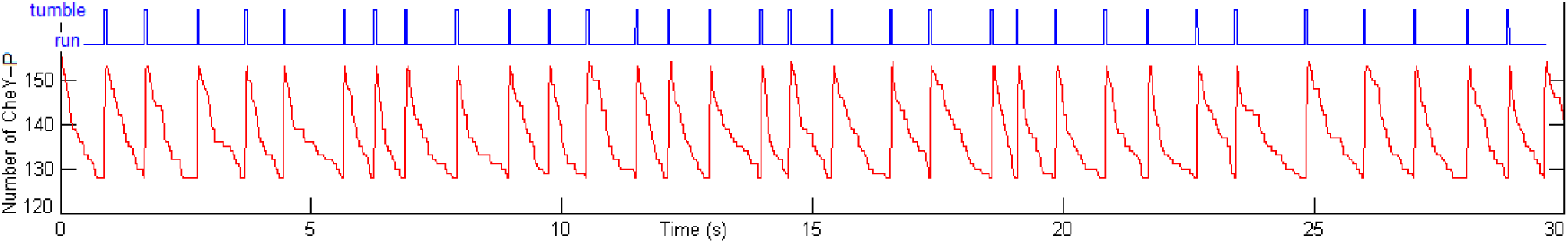
Upper part (blue): Timings of the directional reversal of a single motor. Lower part (red): Almost-periodic variation of the number of CheY-P in the cytoplasm. Looks like the C-ring would "breathe" 24 signal proteins. Due to the small number ofparticles, the statistical fluctuations cause remarkably large time differences until the C-ring has accumulated a sufficient number of CheY-Ps and switches its state abruptly.

**Fig. 4:**
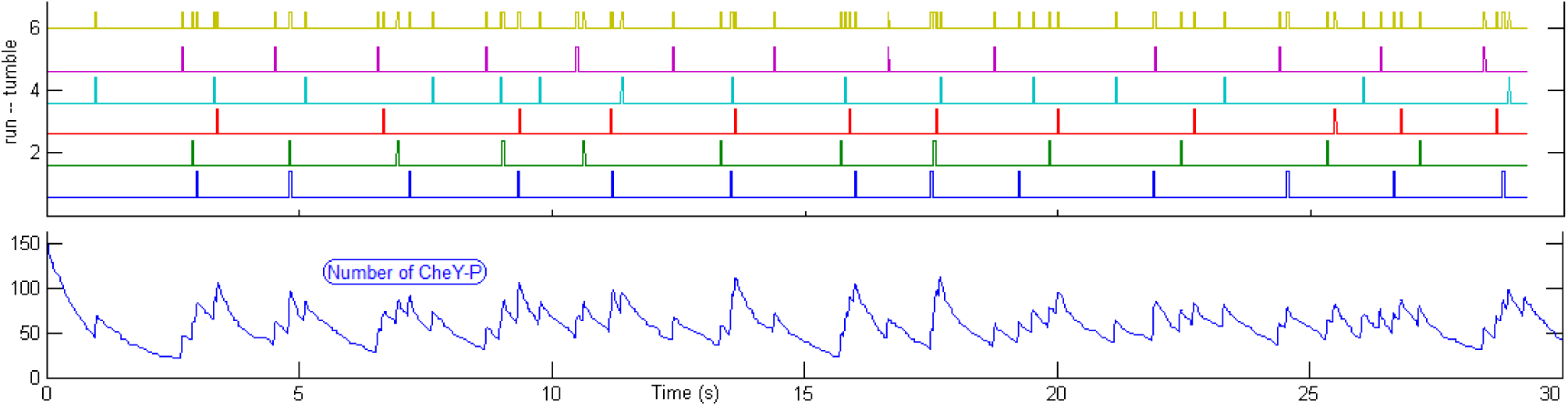
Upper part: Simulated time-dependent rotation direction of five motors. The top curve (gold colored) shows when at least one motor tumbles. Since the cytoplasm contains only 156 CheY-P, it takes longer than in Fig. 3 until the C-rings of the five motors are loaded. The direction of rotation is correspondingly less frequently changed. Below: the number of unbound CheY-P in the cytoplasm varies greatly because the absorption by the motors depends on the direction of rotation. Note the synchronous events at the time points 14 s, 16 s and 18 s, framed by “runs”. Starting the simulation with another random distribution of the CheY-P, the synchrony can be observed more frequently or less frequently. It never disappears.

Strikingly often, several motors tumble almost simultaneously. Then, the number of unbound CheY-Ps in the cytoplasm increases rapidly, and this oversupply accelerates the switching of further motors. This is followed by a longer phase, while the C-rings are reloaded. During this period, E. coli swims straight ahead. It is probably not easy to model these irregular fluctuations with a set of differential equations.

The synchronous reversal of the direction of rotation of several motors follows from the fact that the cytoplasm contains hardly more CheY-Ps than the motors can bind.

## Technical Details

Whether the simulation of a real process is time-consuming depends not only on the underlying physical formulas and the number of particles to be considered. The calculation time can be reduced by means of advantageous programming.

E. coli contains 2.8 million proteins, 20,000 of which are signal proteins whose path has to be followed. Since chemical reactions (= signal transfer) can occur when two suitable molecules meet, the positions must be compared in pairs. Normal programming would be too time-consuming with 400 million comparisons. In order to achieve an acceptable working speed, this value has to be decreased.

It is not necessary to compare molecules that are far away from each other. Therefore, it is useful to sort the data before each comparison. This is facilitated when the three (x, y, z) coordinates of each protein are packed and then processed as a single data unit. If the spatial resolution in the digital 3D model is limited to several hundred steps per coordinate direction, the complete coordinates of each particle inside a cell can be stored in a single 32-bit data word. In our model, the three coordinates are placed as follows in four contiguous bytes.

0xxx xxxx xxxx 0yyy yyyy yy0z zzzz zzzz

In a common computer, a 32-bit data unit is processed very fast with basic integer instructions - much faster than three times as many floating-point arithmetic instructions. This discretization means a maximum of 2047 steps in the x-direction and up to 511 steps in the y‐ and z-directions. With the selected spatial resolution of 3.8 nm, the C-ring of a motor can be described with sufficient accuracy. An exaggerated resolution (smaller boxes) leads to a smaller time step and extended processing time.

The simulation program can be summarized as a pseudocode:

~~~
While time < 30 seconds
    Brownian motion of all CheY-P
    compare the coordinates of all CheY-Ps and all FliMs
    if coincidence CheY-P = FliM is detected AND this FliM has *not* already bound a CheY-P:
       bind CheY-P, remove it from the cytoplasm
    repeat for all five motors:
       if a motor runs AND has bound more than 28 CheY-P:
          switch to tumble,
          kick off the CheY-P into the cytoplasm.
       if a motor tumbles AND if the motor has less than 4 bound CheY-P:
          switch to run
          allow each FliM to bind one CheY-P
    for each time step:
       store the direction of rotation of each motor and the number of unbound CheY-P
~~~

On a laptop, the simulation of a 30 second period real time requires about 90 seconds.

## References

[1] A. Bren and M. Eisenbach, Changing the Direction of Flagellar Rotation in Bacteria by Modulating the Ratio between the Rotational States of the Switch Protein FliM, J. Mol. Biol. (2001) 312, 699–709

[2] T. Duke, N. Le Novere, D. Bray, Conformational Spread in a Ring of Proteins: A Stochastic Approach to Allostery, J. Mol. Biol. (2001) 308, 541–553

[3] S. van Albada, S. Tanase-Nicola, P. ten Wolde, The switching dynamics of the bacterial flagellar motor, Molecular Systems Biology 2009, 5–316

[4] T. Duke, N. Le Novere, D. Bray, Conformational Spread in a Ring of Proteins: A Stochastic Approach to Allostery, Journal of Molecular Biology 308(3):541–53, 2001

[5] P. Lele, H. Berg, Switching of Bacterial Flagellar Motors Triggered by Mutant FliG, Biophysical Journal 2015, Vol. 108, p. 1275–1280

[6] H. Fukuoka, in A Tale of Two Machines_A review of the BLAST meeting, 2013, Mol Microbiol. 2014, 91(1), p. 6–25

[7] Y. Sowa, R. Berry, Bacterial flagellar motor, Quarterly Reviews of Biophysics 41, 2(2008), pp. 103–132

[8] V. Sourjik, Receptor clustering and signal processing in E. coli chemotaxis, Trends in Microbiology, Vol.12 No.12, 2004

[9] P. Cluzel, M. Surette, S. Leibler, An Ultrasensitive Bacterial Motor Revealed by Monitoring Signaling Proteins in Single Cells, Sciencemag, Vol. 287, p 1652–1655, 2000

[10] J. Yuan, H. Berg, Ultrasensitivity of an adaptive bacterial motor, J Mol Biol 2013 May 27; 425(10)

[11] J. Yuan, R. Branch, B. Hosu, H. Berg, Adaptation at the output of the chemotaxis signalling pathway, Nature, 484(7393): 233–236

[12] A. Ishihara, J. Segall, S. Block, H. Berg, Coordination of Flagella on Filamentous Cells of Escherichia coli, JB 1983, p. 228–237

[13] S. Terasawa, H. Fukuoka, Y. Inoue, T. Sagawa, H. Takahashi, A. Ishijima, Coordinated Reversal of Flagellar Motors on a Single Escherichia coli Cell, Biophysical Journal Volume 100, Issue 9, 2011, Pages 2193–2200

[14] H. Fukuoka, Y. Inoue, A. Ishijima, Coordinated regulation of multiple flagellar motors by the Escherichia coli chemotaxis system, Biophysics Vol. 8, pp.59–66 (2012)

[15] M. Sneddon, W. Pontius, T. Emonet, Stochastic coordination of multiple actuators reduces latency and improves hemotactic response in bacteria, PNAS 2012 vol. 109, no. 3, p. 805–810

[16] Oliveira, H. M. and Melo, L. V. Huygens synchronization of two clocks. Sci. Rep. 5, 11548; doi: 10.1038/srep11548 (2015)

[17] Bo Hu, Y. Tu, Coordinated Switching of Bacterial Flagellar Motors: Evidence for Direct Motor-Motor Coupling?, Phys. Rev. Lett. 110, 158703, 2013

[18] M. Li, G. Hazelbauer, Cellular Stoichiometry of the Components of the Chemotaxis Signaling Complex, JB Vol. 186, No. 1, 2004, p. 3687–3694

[19] Y. Sowa, R. Berry, Bacterial flagellar motor, Quarterly Reviews of Biophysics 41, 2(2008), pp. 103–132

[20] A. Wu, T. Davison, C. Jacob, A 3D Multiscale Model of Chemotaxis in Bacteria, 2015

[21] H. Weidner, Tracking the Diffusion of Signal Proteins in Escherichia Coli, 2017

[22] R. Milo, R. Phillips, Cell Biology by the numbers, Garland science, 2015

[23] M. Elowitz et. al., Protein Mobility in the Cytoplasm of B. subtilis, J. OF BACTERIOLOGY, Jan. 1999, p. 197–203

[24] A. Nenninger, G. Mastroianni, C. Mullineaux, Size Dependence of Protein Diffusion in the Cytoplasm of Escherichia coli, JB 2010, Vol. 192, No. 18, p. 4535–4540

[25] BioNumber Details Page: Diffusion coefficient of CheY-GFP 2012

[26] U. Alon, L. Camarena, M. Surette, B. Aguera y Arcas, Y. Liu, S. Leibler, J. Stock, Response regulator output in bacterial chemotaxis, EMBO Journal, Vol.17 No.15 pp.4238–4248, 1998

